# BspA and Pmp proteins of *Trichomonas vaginalis* mediate adherence to host cells

**DOI:** 10.1101/454884

**Authors:** Maria R. Handrich, Sriram G. Garg, Ewen W. Sommerville, Robert P. Hirt, Sven B. Gould

## Abstract

*Trichomonas vaginalis* is one of the most widespread, sexually transmitted pathogens. The infection involves a morphological switch from a free-swimming pyriform trophozoite to an amoeboid cell upon adhesion to host epithelial cells. While details on how the switch is induced and to what proteins of the host surface the parasite adheres remain poorly characterized, several surface proteins of the parasite itself have been identified as potential candidates. Among those are two expanded protein families that harbor domains that share similarity to functionally investigated surface proteins of prokaryotic oral pathogens; these are the BspA proteins of Bacteroidales and Spirochaetales, and the Pmp proteins of Chlamydiales. We sequenced the transcriptomes of five Trichomonads and screened for the presence of BspA and Pmp domain-containing proteins and tested the ability of individual *T. vaginalis* candidates to mediate adhesion. Here we demonstrate that (i) BspA and Pmp domain-containing proteins are specifically expanded in *T. vaginalis* in comparison to other Trichomonads, and that (ii) individual proteins of both families have the ability to increase adhesion performance in a non-virulent *T. vaginalis* strain and *Tetratrichomonas gallinarum*, a parasite usually known to infect birds but not humans. Our results initiate the functional characterization of these two broadly distributed protein families, whose origin we trace back to the origin of Trichomonads themselves.

## Introduction

*Trichomonas vaginalis* is a microaerophilic parasite and member of the Parabasalid family (Paget & Lloyd, 1990), which thrives in the human urogenital tract (Kusdian & Gould, 2014; Mielczarek & Blaszkowa, 2016). The Parabasalids are members of the eukaryotic supergroup Excavata, which includes important parasites/pathogens such as *Leishmania*, *Trypanosoma* and *Giardia* along with many free-living or also mutualistic species (Adl et al., 2005). With more than 270 million new infections occurring annually (WHO 2012; Sutton et al., 2007), *T. vaginalis* infections are very common. In contrast to the Malaria agent *Plasmodium falciparum* for instance, infection prevalence is neither restricted to certain geographical regions, nor dependent on the developmental status of a country but it is associated with people living in resource limited settings (Hirt & Sherrard, 2015; Kissinger, 2015). While the infection in males often remains asymptomatic, it can lead to urethral inflammation and discharge or dysuria. However, in females it is often more pronounced and characterized by vaginal irritation, inflammation and malodorous discharge (Hirt & Sherrard, 2015; Kissinger, 2015). Only a minority of infections, however, lead to fully-developed trichomoniasis (Hirt & Sherrard, 2015; Kissinger, 2015). Given that most *T. vaginalis* infections remain unnoticed, this poses a problem since asymptomatic infections can still elevate the risk of developing cancer, facilitate the acquisition and transmission of HIV and other viruses and are associated with a number of adverse pregnancy outcomes (Petrin et al., 1998; Twu et al., 2014; Hirt & Sherrard, 2015; Kissinger, 2015). Treatments with a 5-nitroimidazole-based derivate are quite effective through the local release of nitro-radicals inside the parasite, the toxin being activated by enzymes of the parasite’s hydrogenosomes (Cudmore et al., 2004; Leitsch et al., 2014). About 10% of *Trichomonas* strains currently diagnosed display some tolerance or even resistance towards metronidazole-based drugs, which appears associated with a reduced expression or complete absence of flavin reductase 1 (Leitsch et al., 2014). Furthermore, patients allergic to metronidazole-based drugs would also benefit from alternative treatment protocols (Kissinger, 2015).

A key characteristic of the infection mechanism of virulent *T. vaginalis* strains is a rapid morphological transition and the secretion of a variety of virulence factors by the parasite. Within minutes upon exposure to host tissue, the free-swimming flagellated cells differentiate into an amoeboid form, a prerequisite for full adherence to human cells (Lal et al., 2006; Noel et al., 2010; Kusdian et al., 2013). The receptors and the subsequent cascade that triggers this shift, or the reason why especially virulent strains display this behaviour, are currently poorly understood. This morphological transition, however, represents a fascinating and excellent example of rapid phenotypic plasticity, which also allows the amoeboid cells to actively migrate across host tissue with the use of the dynamic actin cytoskeleton (Kusdian et al., 2013). Once amoeboid, *T. vaginalis* scavenges host cell substrate, maybe through a mechanism similar to trogocytosis previously described for *Entamoeba histolytica* (Ralston et al., 2014).

*Trichomonas* pathogenesis also involves the secretion of cysteine proteases that degrade the extracellular matrix and other substrates (Sommer et al., 2005; Hernandez et al., 2014), and the secretion of exosomes that can fuse with human cells to deliver their content, which primes host tissue and increases the ability of the parasite to adhere (Twu et al., 2013). It has been speculated that pathogenicity-relevant gene families, such as cysteine proteases and proteins of the Rab family of small GTPases, are specifically expanded in *T. vaginalis* (Carlton et al., 2007). These gene families appear selectively, and to a degree jointly, expressed upon different environmental stimuli (Gould et al., 2013).

Ranging between 158 and 166 megabase pairs, the genome of *T. vaginalis* (strain G3) still remains the largest protist genome so far sequenced (Carlton et al., 2007; Zubacova et al., 2008). This contradicts the usual trend, as parasite genomes tend to shrink as a consequence of evolutionary reduction (Hupalo et al., 2015). The *T. vaginalis* genome is about 7-times the size of the *Plasmodium falciparum* genome and more than 60-times that of the highly reduced *Encephalitozoon cuniculi* genome (Katinka et al., 2001; Zubacova et al., 2008) and also substantially larger than other extracellular parasite such as *Giardia lamblia* (about 14-times) and *Entamoeba histolytica* (about 7-times) (Hupalo et al., 2015). On six chromosomes, *T. vaginalis* encodes a minimum of 46,000 and maybe up to 60,000 proteins (Smith & Johnson, 2011), but with only 65 genes harbouring introns (Carlton et al., 2007). The current *T. vaginalis* genome sequence data is made of a loose collection of around 17,000 individual scaffolds, whose assembly is hindered by the presence of repetitive sequences that represent over 60% of the entire genome (Carlton et al., 2007; Pritham et al., 2007). Both genome and individual gene duplications are thought to cause the expansion of many gene families. The genome is furthermore characterized by an unusual high number of diverse transposable elements. This includes many giant Maverick elements and together they cover about one third of the *T. vaginalis* genome, which might mediate or facilitate genome and/or local duplications (Feschotte & Pritham, 2005; Pritham et al., 2007; Bradic et al., 2014). Among the expanded gene families in *T. vaginalis* are a variety of surface proteins thought to mediate binding of the parasite to host cells and other mucosal commensals of the local microbiota (Hirt et al., 2007; Carlton et al., 2007; Hirt et al., 2013; Bär et al., 2015).

The interaction of parasite surface molecules with the host cell surface is barely understood, although known to be a crucial component that initiates and maintains the infection (Hernandez-Gutierrez et al., 2004; Ryan et al., 2011). The only human binding partner identified so far is galectin-1 that is bound by the lipophosphoglycan (LPG) coat, which covers the parasite’s surface (Okumura et al., 2008), but even that single known interaction was partly challenged (Chatterjee et al., 2015). The palmitoylation of proteins is crucial for adherence, too (Nievas et al., 2018). Early screening of the genome for potential surface proteins unearthed several candidate families (Hirt et al., 2007), including surface proteases such as those of the GP63- and subtilisin family and many proteins with unexplored function, but with homology to infection-relevant surface proteins of prokaryotic and other eukaryotic pathogens. Subsequent proteomic analyses of the *T. vaginalis* surface identified about 140 membrane-bound surface proteins, including several members of the BspA- (Bacteroides surface protein A) and Pmp-(polymorphic membrane protein) family (de Miguel et al., 2010).

Both BspAs and Pmps are surface adhesion proteins of Bacteroidales (FCB group; Bacteroidetes/Chlorobi group; Bacteroidetes; Bacteroidia; Bacteroidales; Tannerellaceae; Tannerella) and Spirochaetales (Spirochaetes; Spirochaetia; Spirochaetales; Spirochaetaceae; Treponema), and Chlamydiales (PVC group; Chlamydiae; Chlamydiia; Chlamydiales; Chlamydiaceae; Chlamydia/Chlamydophila group; Chlamydia), respectively. BspA proteins are leucine-rich repeat (LRR)-containing proteins characterized by a 23 amino acid long repetitive motif (called *Tp*LRR) that is typically found in the N-terminal region (Kobe & Deisenhofer, 1994; Sharma, 2010; Kobe & Kajava, 2001). These surface proteins mediate host-pathogen interactions and promote cell aggregations (Inagaki et al., 2006; Sharma et al., 2005a; Sharma et al., 2005b; Sharma, 2010). Moreover, BspA-deficient mutants of *Tannerella forsythia* are less likely to induce alveolar bone loss in mice (Sharma et al., 2005a), pointing to a pivotal role in virulence. Pmps are chlamydial surface proteins that mediate the initial binding of the obligate intracellular pathogen and eventual invasion into the host cell (Mölleken et al., 2010; Mölleken et al., 2013). Two consecutive tetra-peptide motifs, FxxN and GGA(I/L/V), are found in a repetitive manner in the N-terminal region of Pmps (Grimwood & Stephens, 1999). At least two copies of these motifs are required to mediate adhesion (Mölleken et al., 2010) and antibodies binding these N-terminal repeat motifs reduce the ability of Chlamydiae to infect by up to 95% (Wehrl et al., 2004).

BspAs have been found in other eukaryotic pathogens such as *Entamoeba*, where they were found to be involved in chemotaxis towards a tumor necrosis factor (Silvestre et al., 2015). It is thought that genes encoding BspA and Pmps proteins were introduced into the genomes of eukaryotic pathogens through horizontal gene transfer (HGT) events (Hirt et al., 2002; Noel et al., 2010), rather than representing convergent evolution or differential loss (Ku et al., 2015). *T. vaginalis* G3 encodes 911 BspA-like proteins and 48 Pmps (Noel et al., 2010; Hirt et al., 2011); expression evidence exists for more than half of them (Gould et al., 2013). A few have been localized to the *Trichomonas* surface (Noel et al., 2010; de Miguel et al., 2010), but no dedicated functional analysis of either the *Tv*BspA or *Tv*Pmp proteins has been carried out, and with the recent exception of *Dientamoeba fragilis* from RNA-Seq data analysis (Barratt et al., 2015), the presence and diversity of BspA-like and Pmp-like genes among other Trichomonad parasites remains unknown.

Here, we performed RNA-Seq on five Trichomonads (*Pentatrichomonas hominis*, *Tetratrichomonas gallinarum*, *Trichomitus batrachorum*, *Trichomonas gallinae*, *Trichomonas tenax*) and compared their expression data with that available for *T. vaginalis* to screen in particular for the presence of BspA and Pmp protein encoding transcripts and to unravel their evolutionary trajectory in Trichomonads. Expression of *T. vaginalis* candidate genes encoding BspA and Pmp proteins in a non-adhesive *T. vaginalis* strain and the galliform and anseriform bird-infecting *Tetratrichomonas gallinarum*, provides evidence for the conserved function of both surface protein families regarding adhesion in prokaryotic and eukaryotic pathogens. The unique expansion, particularly striking for the BspA family in *T. vaginalis*, underscores their importance regarding human-specific pathogenicity. Their diversity, for instance the partial absence of membrane-spanning regions and secretory signals in general, might suggest either unknown means of cell surface anchoring (Hirt et al., 2011), divergent version of signal peptides not recognized *in silico*, or functions other than surface-associated adhesion in Trichomonads.

## Results

### RNA-Seq on five Trichomonads

Compared to most of the other Trichomonads, *T. vaginalis* possesses a remarkable big genome (Drmota et al., 1997; Carlton et al., 2007; Zubacova et al., 2008; Smith & Johnson, 2011), raising the question of whether the significant larger gene families observed in *T. vaginalis* correlate with genome size. We were first required to generate RNA-Seq data for Trichomonad parasites and chose five species with a broader phylogenetic distribution and that infect a variety of different hosts (Table 1). Namely, these were *Pentatrichomonas hominis*, *Tetratrichomonas gallinarum*, *Trichomitus batrachorum*, *Trichomonas gallinae* and *Trichomonas tenax*. For functional classification and comparison to the human parasite, the assembled transcriptomes (i.e. the assembled open reading frames, ORFs) were individually blasted against *T. vaginalis*, for which comparable transcriptome data was generated previously (Gould et al., 2013) and a sequenced genome is available (Carlton et al., 2007). Both the number of assembled ORFs among the different analyzed species and the number of ORFs with homologs in *T. vaginalis* differed (Table 1). With 57%, the lowest number of homologs in *T. vaginalis* were found for *Tri. batrachorum* and with 89% the highest number of homologs were found for both *T. gallinae* and *T. tenax* in line with their phylogenetic relationship (Maritz et al., 2014; Kellerová & Tachezy, 2017). Considering the predicted genome sizes of the analyzed Trichomonads (Zubacova et al., 2008), there is no apparent correlation between genome size and the number of expressed genes. This data adds additional credit to the dynamic and expanded nature of the genomes this group of protists is known for (Barratt et al., 2016; Gould et al., 2013).

**Table 1:** RNA-Seq details of the Trichomonads sequenced.

| Species | Strain | Host | Protein Seqs. | Mapping with TrichDB1.3 | Percentage |
| --- | --- | --- | --- | --- | --- |
| *T. gallinae* | GCB | Parasite – intestine galliform and anseriform | 21996 | 1953 | 89 % |
| *T. tenax* | HS-4 | Commensal – oral cavity human | 24071 | 2143 | 89 % |
| *T. gallinarum* | M3 | Parasite – digestive tract pigeon and raptors | 37740 | 2613 | 69 % |
| *T. batrachorum* | BUB | Parasite – intestine amphibian | 34415 | 1950 | 57 % |
| *P. hominis* | PhGII | Commensal – intestine mammals including human | 46458 | 3234 | 70 % |

For comparative analysis, we first screened for those gene families that had already been in the focus of the *T. vaginalis* genome analysis (Carlton et al., 2007) to uncover to what degree they might vary among the different Trichomonads and whether *T. vaginalis* is in any way special. For most cases, only marginal differences were found among the different gene families, but the pathogen of the human urogenital tract did indeed stand out (Fig. 1). On average, the screened transcribed gene families of *T. vaginalis* were about twice the size in comparison to those of the other Trichomonads. In particular genes encoding either for proteins of the BspA or Pmp family stood out. Notably, a similar BspA gene family expansion was observed for *Tri. batrachorum* (Fig. 1), a parasite of the amphibian intestine. Comparing the estimated genome sizes of Trichomonads and the assembled transcriptomes, no direct correlation can be made. Most importantly, it is evident that variants of both the BspA and Pmp gene family, suspected to be associated with host cell adhesion, are expressed by all Trichomonds that we analyzed. However, both families are noticeably expanded only in *T. vaginalis*.

**Figure 1:**
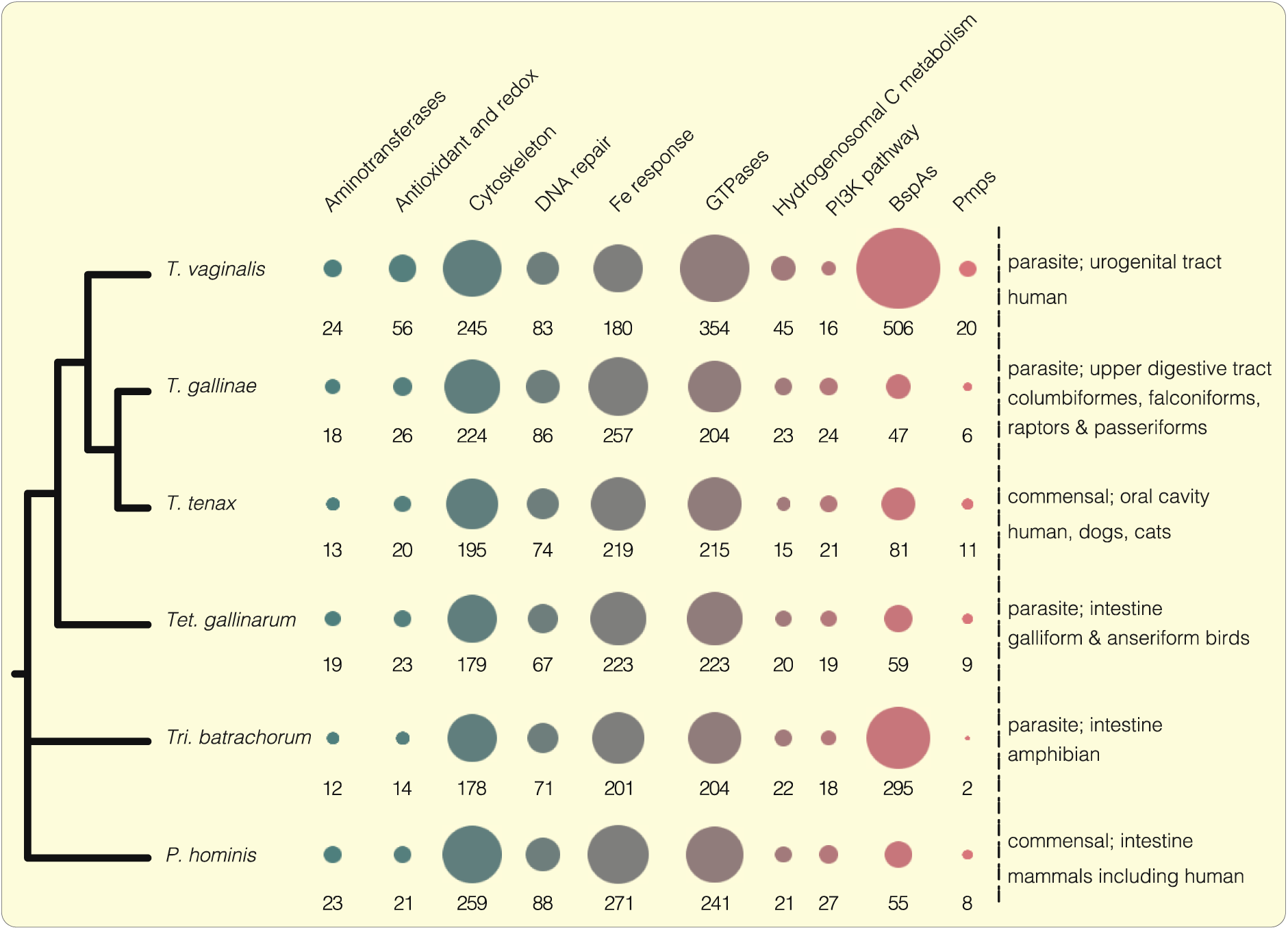
Comparative analysis of specific gene family sizes among Trichomonadida. For our analysis we selected different Trichomonadida species, both commensal and parasitic ones, with a high diversity of host cells. The amount of proteins belonging to different housekeeping gene families or proteins possessing BspA and Pmp specific domains are represented as bubbles. On average the family sizes of *T. vaginalis* are doubled compared to the other Trichomonadida. Especially those genes involved in iron response mechanisms or belonging to the BspA as well as Pmp protein family are almost exclusively expanded in *T. vaginalis*.

### The BspA and Pmp protein family of Trichomonads

The expansion of both the BspA and Pmp protein family in the human pathogen *T. vaginalis* raises the question about their distinct roles during infection, or more precisely whether those proteins are implicated in directly mediating adhesion to human host tissue as has been previously speculated (Hirt et al., 2007; Noel et al., 2010). Structural comparison of the two families reveals several similarities among the Trichomonads, which are consistent with the built up found for prokaryotic homologs, but with a few important exceptions (Fig. 2). Similar to their prokaryotic counterparts, the Pmp-specific repeat motifs FxxN and GGA[I/L/V], and the leucine-rich-repeats of the BspA family are present throughout the main parts of the respective proteins towards their N-termini (Grimwood & Stephens, 1999; Sharma et al., 1998). Besides these conserved regions, both families have experienced similar modification towards their C-terminus, in particular the substitution of prokaryote-specific elements (such as the por secretion system or the autotransporter domain) (Dautin & Bernstein, 2007; Shoji, 2011) with a single membrane spanning domain close to the C-terminus.

**Figure 2:**
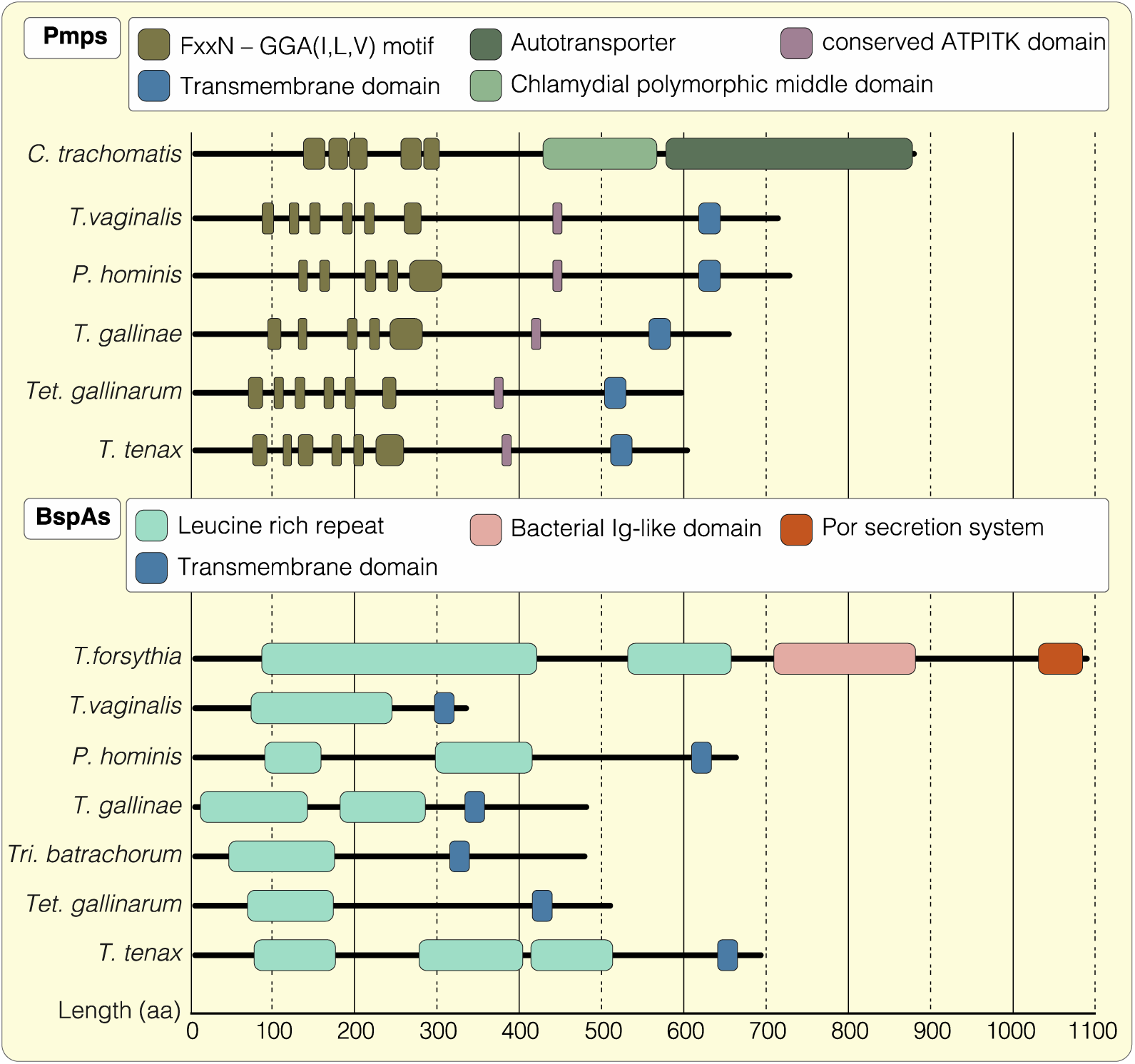
Structural comparisons of the Pmp and BspA protein family in different Trichomonadida. *(top)* Schematic illustration of selected Pmp structures in different Trichomonadida together with the prokaryotic model *Chlamydia trachomatis*. (Accession numbers: *C. trachomatis*, gi 34539119; *T. vaginalis*, TVAG_249300; *P. hominis*, PEHa011017; *T. gallinae*, TEGb004672; *Tet. gallinarum*, TRGa004464; *T. tenax*, TRTa003481). The Pmp family is characterized by multiple repeats of the FxxN and GGA[I/L/V] motifs located in the N-terminal region but compared to *C. trachomatis* the Trichomonad proteins miss the C-terminal polymorphic middle and autotransporter domain. Instead they possess a conserved ATPITK motif as well as a transmembrane domain at their C-terminus. *(bottom)* Schematic illustration of the protein structures of selected members the BspA family in different Trichomonadida compared to *Tannerella forsythia*. (Accession numbers: *T. forsythia*, gi 3005673; *T. vaginalis*, TVAG_240680; *P. hominis*, PEHa029834; *T. gallinae,* TEGb004448; *Tri. batrachorum*, TRBa028008; *Tet. gallinarum*, TRGa003876; *T. tenax*, TRTa008806). The BspA protein family is unified by several N-terminal copies of leucine-rich-repeat elements. In the *T. forsythia* the BspA additionally possess a bacterial Ig-like domain (Big-domain) as well as a por secretion system both localized towards the C-terminus. In the Trichomonads those are replaced by a transmembrane domain in a proportion of BspA-like proteins.

*T. vaginalis* is an extracellular parasite. During infection it faces host immune defence mechanisms and interacts with the urogenital microbiota, of which some is also phagocytosed (Juliano et al., 1991; Pereira-Neves & Benchimol, 2007). The abundance of BspA and Pmp proteins with endocytic motifs likely reflects a specific need and perhaps an adaptation to the host habitat; the question follows whether the endocytic machinery was expanded in a similar way. We screened for proteins required for vesicle formation at the plasma membrane (Pearse, 1975; Keen, 1985; Takei & Haucke, 2001), vesicle fusion (Pelham, 1995, Jahn & Scheller, 2006), as well as proteins involved in intracellular trafficking (Seaman, 1998; Lee et al., 2004) such as those that mediate endosome/Golgi and Golgi/ER transport. Indeed, in comparison to another anaerobic human parasite, *Giardia intestinalis*, the protein families in question were found to be expanded among all Trichomonads and especially so in *T. vaginalis* (Fig. 3A). The Rab subfamily of small GTPases stood out in particular. Representing 75% of the GTPases in the parasite (a single celled organism), the contribution of the Rab family is remarkably higher than in humans (a metazoan with tissue-specific expression), where they comprise less than 50% in total (Colicelli, 2004). This is comparable to the *T. vaginalis* BspA and Pmp family, which is also specifically expanded. Hence, it seems likely that those proteins are involved in the endocytic machinery, too.

**Figure 3:**
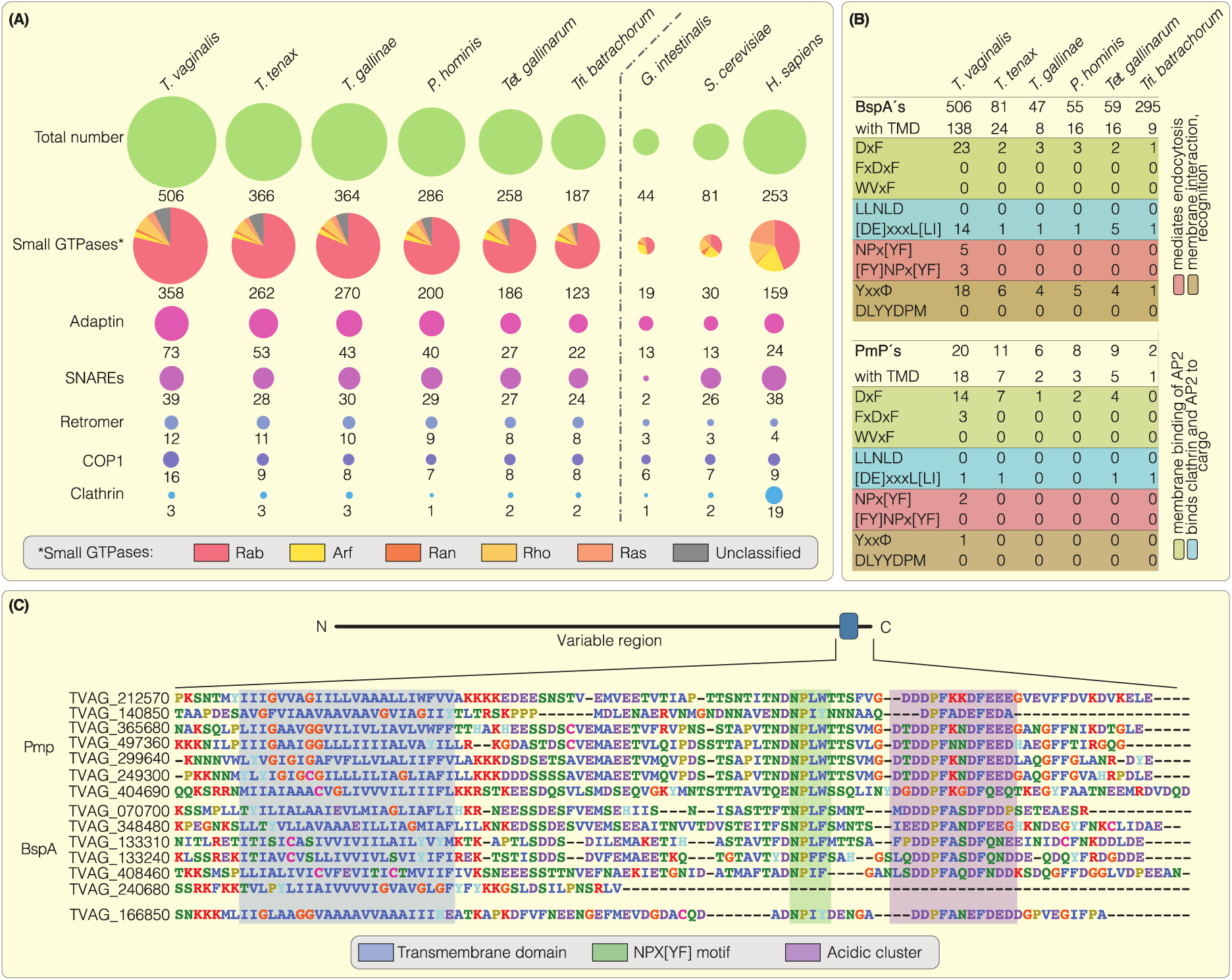
Expansion of endocytosis-related protein families and analysis of specific endocytic motifs within the Pmp and BspA protein family. *(A)* The sizes of several protein families involved in endocytosis are represented as bubbles and compared among the Trichomonadida. Although *T. vaginalis* again shows the highest numbers, all gene families are expanded in the Trichomonads compared to *Giardia intestinalis* another anaerobic human parasite. In particular the Rab subfamily of the small GTPases displays a specific expansion in the Trichomonadida as obvious by comparing those numbers to the human gene family. *(B)* The Trichomonadida Pmp and BspA families were screened for the presence of a transmembrane domain and further for specific endocytic motifs within their cytoplasmic tails. While 90% of the *T. vaginalis* Pmps carry a TMD it is just around a quarter for the BspAs. Notably, in *Tri. batrachorum* which showed a similar expansion of this protein family only around 3% exhibit a TMD. The fraction of proteins with motifs involved in endocytic processes is divers. However, the *T. vaginalis* proteins possess an increased amount of specific motifs e.g. the NPx[YF]. *(C)* Sequence alignment of *T. vaginalis* Pmp and BspA proteins with focus on the C-terminal region. Together with the conserved transmembrane domain those proteins – with one exception (TVAG_240680) – also carry the *T. vaginalis* specific NPx[YWF] motif and an acidic cluster within their cytoplasmic tails.

The initial screen for proteins possessing a TMD generally resulted in comparable relative values. Only in *Tri. batrachorum*, which showed an expansion of the BspA family similar to that of *T. vaginalis*, the amount of TMD containing proteins is considerably lower. Screening these proteins for motifs uncovered several that are recognized by the endocytic machinery (Fig. 3B), especially located within the C-terminal tails. We found a DxF motif involved in membrane binding of the AP2 complex (Bonifacino & Traub, 2003; McMahon & Mills, 2004) present in proteins of almost all Trichomonadida analyzed, while the NPx[YF] motif seems to be restricted to *T. vaginalis*. Sequence alignment of selected Pmps revealed a frequent modification of this motif, in particular a substitution of phenylalanine with tryptophan in *T. vaginalis*. This substitution does not necessarily render the motif dysfunctional. Similar functional substitutions have been observed, such as for a targeting motif of secondary red algae (Gruber et al., 2007). Furthermore, the presence of a conserved acidic cluster in both protein families is evident (Fig. 3C), which represents another family of membrane sorting signals (Bonifacino & Traub, 2003; Navarro Negredo et al., 2017). Nevertheless, there is a large diversity among both protein families with regard to the presence or absence of domains in a single protein and overall no generalized pattern is recognizable.

### Individual *T. vaginalis* BspA and Pmp proteins mediate adhesion

In order to investigate the influence of the Pmp and BspAs on adhesion, we selected candidate proteins and expressed them in the non-adhesive *T. vaginalis* T1 strain and the bird pathogen *Tetratrichomonas gallinarum*. We chose candidate proteins (BspA TVAG_240680, Pmp TVAG_212570 and Pmp TVAG_140850), that are generally expressed at higher levels or even upregulated upon exposure to host cells (Gould et al. 2013), and which were identified through cell surface proteomics on the pathogen's surface and displayed an increased abundance in highly adherent strains (de Miguel et al., 2010).

T1 clones expressing candidate proteins were used to perform adhesion assays on a monolayer of vaginal epithelial cells (VECs). All tested candidate proteins increase the adherence of *T. vaginalis* to VECs two to four-fold in comparison to the T1 wt strain (Fig. 4A). This is only half of the number of adhering cells counted for the highly virulent FMV1 strain, but still matches the results of the positive control (TVAG_166850) that was previously shown to facilitate the adhesion of *T. vaginalis* (de Miguel et al., 2010). The expression of malic enzyme (TVAG_183790), a protein of hydrogenosomal energy metabolism (and our negative control), did not lead to increased adherence and delivered results comparable the T1 wildtype. Furthermore, we expressed the BspA and one Pmp candidate protein in *Tet. gallinarum*, a parasite usually infecting the digestive tract of birds, and analyzed their influence on the binding to vaginal epithelial cells. Although the overall adherence was lower compared to *T. vaginalis* clones expressing the same candidate proteins, both lead to a significant increase of adhering parasites. Compared to the M3 wildtype strain, the number of adhering cells was more than 1.5-fold increased in the case of the Pmp, and more than 4-fold higher for the *Tet. gallinarum* clone expressing the BspA protein (Fig. 4B).

**Figure 4:**
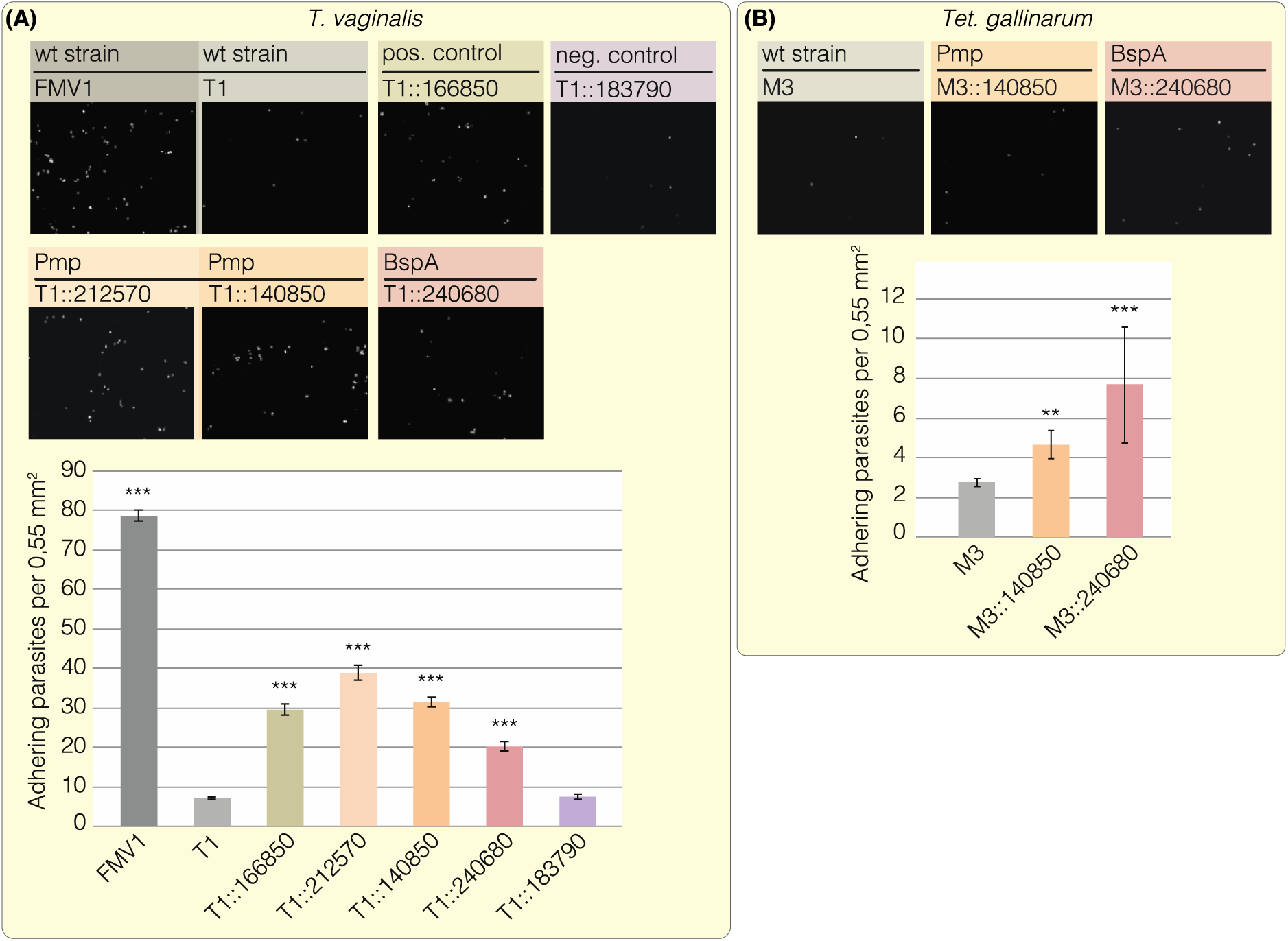
Adhesion assays for *T. vaginalis* and *Tet. gallinarum* on vaginal epithelial cells. *(A)* Overexpression of specific Pmp and BspA proteins increases the adherence of the non-infective T1 strain to vaginal epithelial cells (VECs). Example pictures for every assay are shown on the top where each white dot represents one adhering parasite. Compared to the T1 wt expression of the candidate Pmp (TVAG_212570 and TVAG_140850) and BspA (TVAG_240680) proteins significantly increased the number of attached *T. vaginalis* cells. The highly infective FMV1 wt strain and malic enzyme (TVAG_183790) were used as a positive and negative control respectively. Adhesion assays were performed three times in biological triplicates and for each candidate a total of 108 pictures were analysed. T-Tests were performed for the analysis of statistical significance compared to *T. vaginalis* wildtype strain T1 (***, P<0,0001; **, P<0,001; *, P<0,05). *(B)* The expression of TVAG_140850 (Pmp) and TVAG_240680 (BspA) also significantly increases the ability of the bird infecting pathogen *Tet. gallinarum* M3 wt to adhere to human host cells. Adhesion assay was performed once in biological triplicates and for each candidate a total of 36 pictures were analysed. T-Tests were performed for the analysis of statistical significance compared to *Tet. gallinarum* wildtype strain M3 (***, P<0,0001; **, P<0,001; *, P<0,05).

### BspA and Pmp proteins localize predominantly to internal compartments, not the plasma membrane

We first analysed the subcellular localization of the candidate proteins used for the adhesion assays. Both the Pmp and the BspA protein, as well as the positive control (TVAG_166850), localize to structures that are reminscent of the endoplasmic reticulum (ER) and Golgi apparatus of the parasite (Fig. 5A) (Burstein et al., 2012; de Andrade Rosa et al., 2014; Riestra et al., 2015). Since most tested proteins were identified as part of the surface proteome of *T. vaginalis* (de Miguel et al., 2010), and one would hence expected them to localize to the parasite’s plasma membrane, we performed a minimum of at least three independent experiments for each protein, but never found the proteins to localize to the surface.

**Figure 5:**
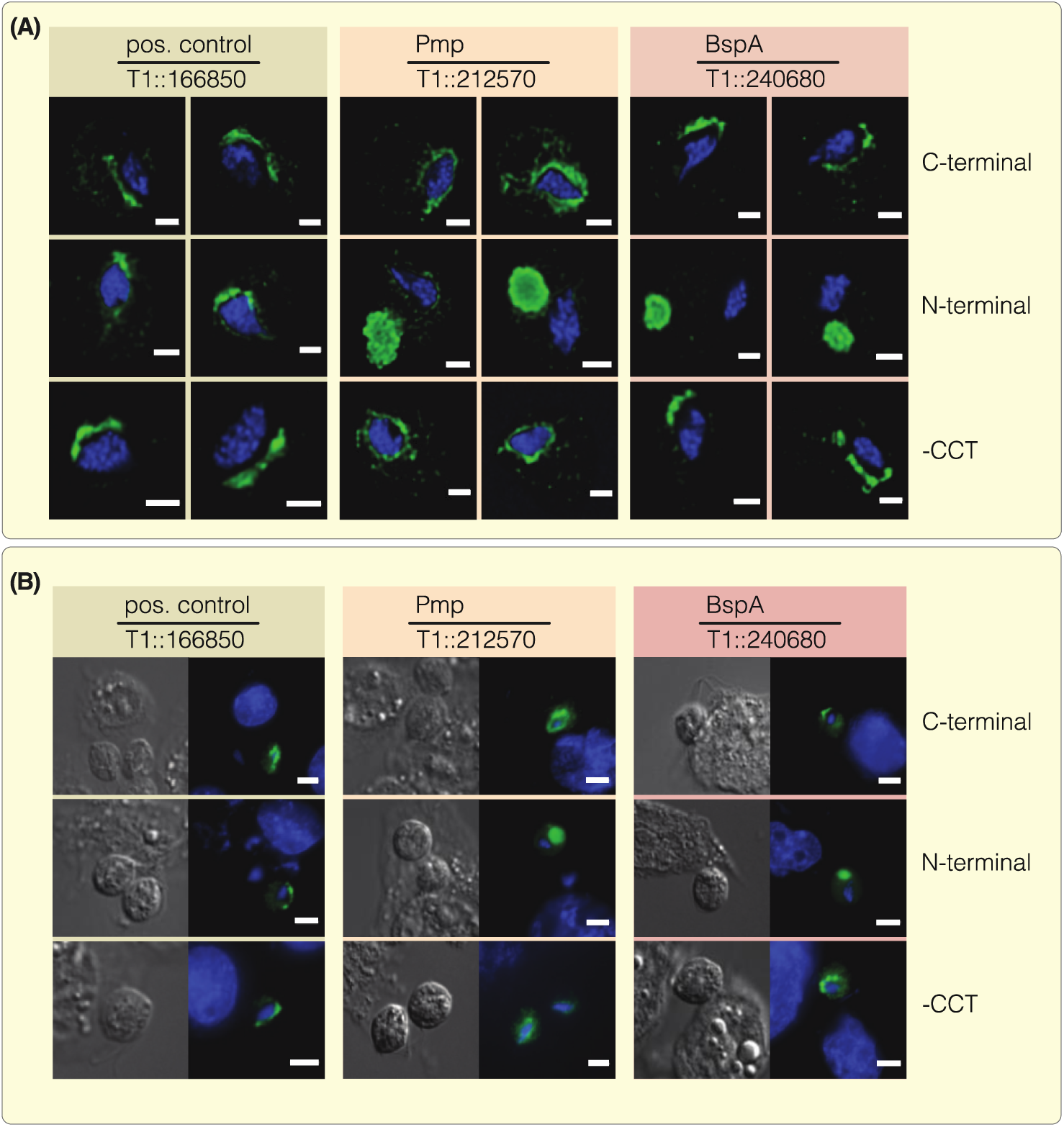
BspA and Pmp proteins localize predominantly to the ER and Golgi. The subcellular localizations of the positive control (TVAG_166850) and the candidate Pmp (TVAG_212570) and BspA (TVAG_240680) protein were analysed using a specific antibody against the HA-tag. The protein localizations are shown (green) together with the nucleus (blue) which was stained using 4′,6-Diamidin-2-phenylindol (DAPI). *(A)* The C-terminal tagged proteins which were used for adhesion assays constantly show a localization to the ER and Golgi apparatus. While shifting this tag position to the N-terminus has no influence on the positive control – which is still found to resides in the ER – it leads to a relocalization of both the Pmp and BspA to the lysosome. In contrast, removing the cytoplasmic tails – by replacing it with the HA-tag – did not lead to any localization shift compared to the C-terminally tagged proteins analyzed first. Scale: 2 µm. *(B)* We further checked if those localizations are influenced by the presence of host cells, but no change could be observed for *T. vaginalis* cells grown on VECs. Scale: 5µm.

To exclude the possibility that the ER and Golgi localization was a result of C-terminal HA-tagging, which could potentially interfere with the also C-terminally localized endocytic motifs, we additionally tagged the proteins at their N-terminus. While this switch had no influence on the localization of the postive control, it changed the localization of the Pmp (TVAG_212570) and BspA (TVAG_240680) protein (Fig. 5A) and the number and size of the lysosomes. Both N-terminally tagged constructs localize to a single spherical large lysosome, which is evident by the colocalization with LysoTracker (Fig. 6). However, in cell lines expressing C-terminally tagged constructs, we observed many small lysosomes (Fig. 6), which corresponds to what is usually described for *T. vaginalis* (Burstein et al., 2012; Huang et al., 2014). The clones in which the fusion proteins localize to the lysomes, also perform poorly in terms of increasing adhesion performace (supplementary figure S1). Hence, the N-terminal tagging of the BspA and Pmp proteins and their incorporation into lysosomes leads to the formation of one single large lysosome, instead of many small ones. In contrast the substitution of the cytoplasmic tails that carry the motifs known to be recognized by the endosomal machinery, through an HA-tag, did not alter ER/Golgi localization (Fig. 5A).

**Figure 6:**
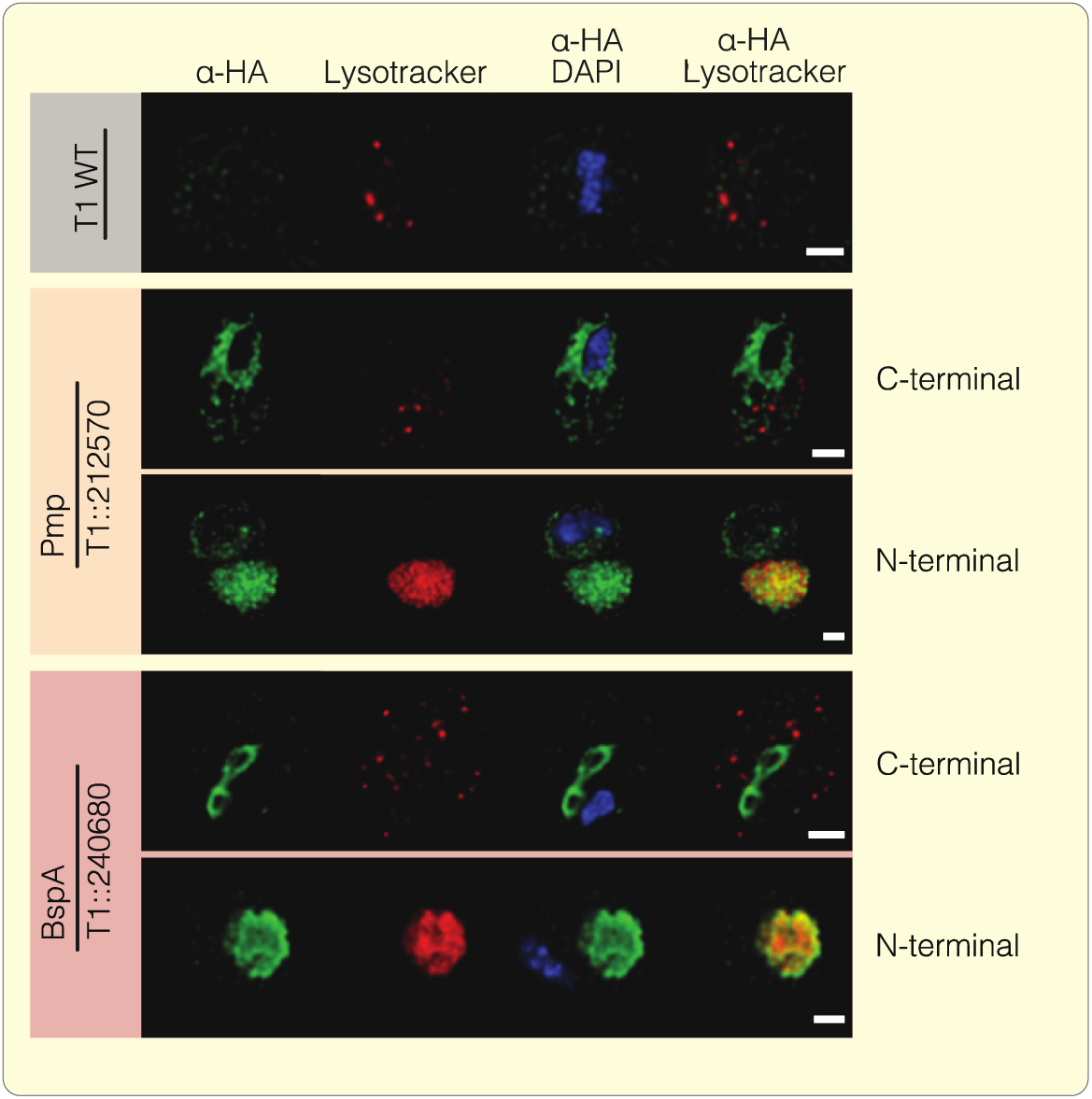
Lysosome localization of the N-terminal tagged Pmp and BspA protein. To verify the observed localizations of the N-terminal tagged Pmp and BspA protein – which occurred to be lysosomal – a colocalization with LysoTracker (red) was performed. The candidate proteins were detected using a specific α-HA antibody (green) while 4′,6-Diamidin-2-phenylindol (DAPI) was used for nucleus staining (blue). In the T1 wt strain as well as the candidate proteins which carry the HA-tag at the C-terminus several small lysosomes were detected, equally distributed in the cytosol. Simultaneously the Pmp and BspA show circular structures around and near the nucleus respectively. In contrast, one greatly enlarged lysosome could be detected in the N-terminally tagged cultures which clearly colocalizes with the HA-fusion constructs. Scale: 2 µm.

We also considered whether the exposure to host tissue, vaginal epithelial cells (VECs), could trigger a change in localization. However, the observed localization of the proteins for cells growing in the absence of VECs did not differ from those that were exposed (and found attached) to VECs (Fig. 5B). Similarly, we expressed two of our candidate proteins, the positive control and the BspA, in the highly infective FMV1 wildtype strain in order to exclude the possibility that the missing ability to fully adhere has an influence on the protein localizations, yet no localization change was observed (supplementary Fig. 2). Furthermore, other detergents were tested to ensure that no over-extraction of membrane-located proteins occurred, which has been observed for protocols using Triton X-100 alone (Sharma et al., 2008). Neither the use of NP-40, nor treatments with the mild detergent digitonin, however, led to the proteins also being observed to localize to the plasma membrane – the localization pattern remained the same as with using Triton X-100 (supplementary Fig. 3). Finally, to circumvent the use of any detergent, we tagged the proteins with the green fluorescent protein for live imaging. While the fluorescence was low (GFP requires O2 to fluoresce, but *Trichomonas* is an anaerobic parasite), again only a localisation to the ER and Golgi was evident (supplementary Fig. 4).

## Discussion

Infections with *T. vaginalis* continue to increase and remain an underestimated threat due to the often asymptomatic progress of infection (Edwards et al., 2016); *T. vaginalis* is apparently more often a commensal than a parasite. Any *T. vaginalis* infection, including asymptomatic ones, is accompanied by small inflammatory reactions of the affected host tissue, which are caused by several kinds of different processes. They include in particular the secretion of a substantial amount of cysteine proteases (Sommer et al., 2005) and lesions caused by the attachment of the parasite to urogenital tract tissue and the phagocytic uptake of its cells (Midlej & Benchimol, 2010). The genome of *T. vaginalis* is characterized by an unusually large number of repetitive elements and transposons and the massive expansion of gene families (Pritham et al., 2007; Carlton et al., 2007; Bradic et al., 2014). It is thought that especially those gene families that encode proteins associated with infection are expanded (Carlton et al., 2007) and expressed in a coordinated manner (Gould et al., 2013).

So far only the genome of *T. vaginalis* has been fully sequenced, but genome size estimations exist for several other Trichomonads (Zubacova et al., 2008). One observation we made is that no clear correlation between the predicted genome sizes and the number of expressed genes is evident. Until genome data for other Trichomonads than *T. vaginalis* become available, such numbers need to be treated with caution. Hence, we decided to only compare the numbers of expressed genes, also regarding *T. vaginalis*. Doing so demonstrates that among the different Trichomonadida considered here, *T. vaginalis* generally expresses the largest number of genes from each family analyzed, but in particular the BspA and Pmp families stand out. These numbers, in particular those of the BspA family, highlight the importance of the two gene families for the parasite and their universal presence among the Trichomonads we sequenced, tells us something about their origin.

Previous phylogenetic analyses have suggested that the BspA and Pmp family were acquired by *T. vaginalis* through the horizontal transfer of DNA (HGT) from different prokaryotic sources (Hirt et al., 2002; Alsmark et al., 2009). RNA-Seq data of all five Trichomonads we sequenced in addition to *T. vaginalis*, contained a substantial number of transcripts encoding proteins of the Pmp and BspA family. This gene distribution argues for an ancient origin in Trichomonads, which is supported by the additional presence of the BspA family in the Tritrichomonad *Dientamoeba fragilis* (Barratt et al., 2015). One could argue for an ancient acquisition from mucosal-dwelling prokaryotic microbiota [although Trichomonad BspAs share the best homology with the eukaryote *Entamoeba dispar* (Noel et al., 2010)], also in terms of the proteins conserved function in prokaryotes and, in this case, the eukaryotic pathogen *T. vaginalis*. A simple blast, however, also identifies BspA-like sequences in some hapytophyte algae (e.g. *Chrysochromulina sp.* KOO53905.1) and the coral *Stylophora* (XP_022807022.1), which complicates interpretations if they are not contaminations. Independent acquisitions through HGT cannot be ruled out, but the reasons for retention will differ from those in Trichomonads.

Horizontal gene transfer in eukaryotes, or more specifically its frequency, remains a controversial topic (Ku et al., 2015; Husnik & McCutcheon, 2018). If these proteins trace back to HGT and a prokaryotic source, we can conclude that it occurred before the diversification of parabasalia. Any phylogenetic analysis is impeded by the low sequence identity of eukaryotic BspAs and Pmps, which is furthermore restricted only to the repetitive motifs of the N-termini that are thought to be essential for the interaction with host tissue cells. Hence, if they are of prokaryotic origin, then early in their parabasalid evolution, the C-terminal prokaryotic domains (e.g. the por secretion system and autotransporter domains) were substituted with eukaryotic ones, which often contained a single TMD and endocytotic motifs (Fig. 2 and 3C). Not entire genes, but only the parts useful for the eukaryotic parasite, namely the extracellular domain containing the *Tp*LRR, were retained and used as building blocks and, especially in *T. vaginalis* and *Tri. batrachorum*, were expanded.

Concomitant with the expansion of the BspA and Pmp protein family in *Trichomonas* and the presence of C-terminal motifs known to be recognized by the endocytic machinery (Fig. 3B), protein families associated with vesicle trafficking are also expanded. For all six Trichomonads, the gene families of the adaptin-, COP-, snare-, retromer and especially small GTPases show a significant expansion in comparison to the anaerobic parasite *G. intestinalis* or yeast (Fig. 3A). In comparison with human, the gene family sizes are comparable if not larger (except for the clathrin family; Fig. 3A), which is even more astonishing considering that *T. vaginalis* expresses all proteins as a single celled organism and does not demonstrate tissue specificity. This expansion underscores the importance of endocytic uptake of extracellular substrate in *T. vaginalis*. Furthermore, it raises the question of whether the combination of an N-terminal domain required for adhesion with a C-terminal domain recognized by the endocytic machinery hints at trogocytosis similar to that observed in *Entamoeba* (Ralston et al., 2014), or perhaps a rapid recycling of the plasma membrane and exchange of surface molecules to evade the human immune system as it occurs in trypanosomes (Batram et al., 2014).

Expression of a BspA (TVAG_240680) as well as Pmp (TVAG_212570 and TVAG_140850) proteins significantly increases the ability of T1 parasite cells to adhere to human tissue (Fig. 4A). Overexpression of the candidate proteins boosted the adhesion capacity of the T1 strain up to almost 50% indicating that those protein families are key factors for infection. Furthermore, the failure of the malic enzyme (TVAG_183790) to do the same, corroborates doubts about a potential moonlighting function of hydrogenosomal proteins in the parasite's adhesion (Addis et al., 1997; Hirt et al., 2007; Twu et al., 2013; Kusdian & Gould, 2014).

Details of how the Pmp and BspA proteins are involved in adhesion and how they might subsequently be recycled remain obscure. Our sequence analysis shows that both members of the Pmp protein family analyzed possess endocytic motifs within their cytoplasmic tails that are essential for clathrin mediated endocytosis, for example through interactions with the AP2 complex (Fig. 3C). Furthermore, our data hints to a putative function in mediating host specificity. Expression of the candidate Pmp TVAG_140850 and BspA TVAG_240680 proteins also significantly increases the ability of *Tet. gallinarum* – a pathogen naturally infecting birds – to adhere to human tissue (Fig. 4B). For chlamydial Pmps it was suggested that the specific FxxN and GGA[I/L/V] motifs are either directly or indirectly involved in mediating interactions with human receptors (Mölleken et al., 2010). Based on this, Pmps possibly recognize specific host cell structures subsequently leading to the endocytosis of host material through trogocytosis.

The LRR domain of BspA proteins possesses putative functions in binding of host cell receptors in *T. forsythia*, likely triggering a signalling cascade that promotes bacterial invasion (Inagaki et al., 2006). *T. vaginalis*, however, is an extracellular parasite, but consistent with its greater impact on the adhesion of *Tet. gallinarum* – a more than 4-fold increase compared to the wildtype – this protein is likely involved in host–parasite interactions through the binding of specific human receptors. Moreover, *T. vaginalis* was shown to secrete exosome-like vesicles, which play a role in the parasite's attachment to host cells and modulate the human immune response (Twu et al., 2013). Since a BspA protein (TVAG_240680, with a TMD and a 15 residues CCT without obvious signals for endocytosis) was found to be present in the pathogen's exosome proteome, it might be associated with priming host cell tissue for parasite binding and eventual colonisation; possibly through *Tv*BspA (on the parasite)-*Tv*BspA (on human cells) interactions as shown for *Tannerella forsythia* and *Treponema denticola* and other bacteria (Sharma 2010). Similarly, some parasite BspA proteins could also contribute to parasite binding to prokaryotes and other members of the microbiota to eventually mediate their phagocytosis (Juliano et al., 1991; Pereira-Neves & Benchimol, 2007).

In prokaryotes, BspA and Pmp proteins aid attachment to host tissue and in Trichomonads, at least for those that carry a TMD, it seems natural to assume that they are anchored into the plasma membrane of the eukaryotic parasite. However, it is evidently more complicated than that. The BspA and Pmp HA-fusion constructs localize predominantly to intracellular compartments, but not the plasma membrane (Fig. 5A, supplementary figures S2-S4). The defined localization around the entire nucleus is typical for the endoplasmic reticulum of the parasite that embraces the nucleus in several layers and is largely absent from the remaining cytosol, while the two adjacent rings at the apical end, and in close proximity to the nucleus, are typical for the Golgi apparatus of Trichomonads (Burstein et al., 2012; Andrade-Rosa et al., 2013). A sole localization to compartments of the endomembrane system appears unlikely, as it is incompatible with the detection of some as part of a surface proteome analysis (de Miguel et al., 2010), the presence of the BspA protein TVAG_240680 in exosomes (Twu et al., 2013), and the increasing adhesion to host tissue we observed.

Still, ER/Golgi localization for TVAG_166850 was observed before and agrees with a former statement, in which expression of the full-length protein led to retention in the ER (Riestra et al., 2015). This targeting to the endosomal compartments occurs independently of N-terminal signal peptides, as none are detectable for any of the proteins we analyzed. Perhaps the majority of the protein resides in the ER and Golgi and is only transported to the surface in small concentration. The TMD of TVAG_166850 was shown to possess a cleavage site recognized by a specific membrane located rhomboid protease (*Tv*ROM1) (Riestra et al., 2015). This finding suggests temporary surface localization of this protein followed by a cleavage-induced secretion, which could provide an explanation for missing membrane localization. However, there is an inconsistency regarding its role for the attachment of the parasite since both increased, as well as an inhibited cleavage by *Tv*ROM1 leads to a higher adherence of *T. vaginalis* (Riestra et al., 2015).

Interestingly, for the Pmp and BspA protein, the N-terminal tag position leads to a localization shift to the lysosome. Moreover, this has a significant impact on lysosome morphology (Fig. 6). Instead of many small lysosomes frequently distributed in the cytosol, cultures expressing N- terminally tagged BspA and Pmp proteins show one lysosome, which is massively expanded. This is possibly due to the high number of incorporated proteins. The switch from a C- to a N-terminal tag also interferes with their inferred function leading to a significant decrease in the ability to mediate adhesion (supplementary Fig.1), which furthermore underscores that the observed, dominant ER/Golgi localization is part of the proteins native localization and function in mediating adhesion. For the Pmp protein this is supported by former findings, where it was shown that Pmp21 from *Chlamydia pneumoniae* is processed post-translational, leaving only the N-terminal part that acts as adhesin and therefore is essential for activity (Wehrl et al., 2004).

The localizations observed might overall complicate interpretations, but they are robust. Neither the use of different detergents nor GFP-tagging (and live-imaging) changed the localizations. Other proteins expected to be anchored into the plasma membrane have been observed to predominantly localize to the secretory system (Riestra et al. 2015; Coceres et al. 2015), suggesting a more common mechanism is behind this in *Trichomonas*, which a dedicated future investigation needs to tackle.

There is a wealth of diversity amongst the Pmp and especially BspA protein families of *T. vaginalis*. The parasite expresses many hundreds of BpsA genes and at least two-dozen Pmp-encoding genes, all of which differ in their sequence and even domain architecture. Only about 27% of the BspA, but roughly 90% of the Pmp proteins we found to be expressed, carry a TMD (Fig. 3A) that would allow them to be anchored into the plasma membrane to serve direct parasite adhesion. The same is true for the motifs recognized by the endocytotic machinery; they are only infrequently found and rarely in combination with the other mentioned domains (Fig. 3C). This diversity can explain the differences we observed in particular in terms of localization behavior. It appears as if these two protein families are part of an ‘evolutionary playground’, with little selection pressure acting on a conserved set of domains simultaneously, with some domains being shared between the two families as well as other candidate surface proteins (shared TMD-CCT with signals for endocytosis, Fig. 1C) suggesting gene fusion events combining a given TMD-CCT with various extracellular domains.

## Conclusion

Our results show that, although present in all Trichomonads analyzed, the massive expansion of both the Pmp and BspA protein family is restricted to *T. vaginalis*. This indicates that those proteins play an important role for the infection of the human host. Furthermore, the common presence among the Trichomonads provides evidence for ancient origin, maybe through HGT, which occurred prior to the early evolution of the parabasalids. The presence of several endocytic motifs within both protein families suggests that some of them play a role either in the endocytosis of host material or possibly the evasion of the host immune system by dynamically remodeling the surface proteome, both of which are essential components of the *T. vaginalis* infection. This is further underlined by the increased number of proteins associated with the endocytic machinery such as proteins associated with vesicle formation and intracellular trafficking. We demonstrated that the expression of specific BspA and Pmp proteins significantly increased the adherence of the non-infective T1 strain of *T. vaginalis*, as well as the ability of the bird infecting *Tet. gallinarum* to bind to human host tissue. The latter confirms their involvement in the adhesion process and hints at a putative role in mediating host specificity. In contrast, the shared cell surface BspA and Pmp protein families across the investigated Trichomonads and the demonstration that some family members are involved in binding to host cells, might have contributed to the zoonotic potential of some of these parasites, assuming that one or more family members bind to shared mucosal features across birds and mammals including human (Maritz et al., 2014). In particular the bird infecting *T. gallinae* closely related to the buccal *T. tenax* from mammals, and now known to be common among both pet mammals and human, might be a particular point in case (Maritz et al., 2014; Kellerová & Tachezy, 2017). What remains puzzling is the localization to the secretory system observed by others (e.g. Riestra et al. 2015; Coceres et al., 2015) and us regarding what one would predict are proteins anchored to the surface. Together with the change in lysosome morphology, and expansion in proteins that orchestrate vesicle biology, this hints at unexplored avenues of *T. vaginalis* molecular cell biology.

## Material and Methods

### Culturing

*Trichomonas vaginalis* strains T1 and FMV1 were cultured in tryptone-yeast extract maltose medium {2,22% (w/v) tryptose, 1,11% (w/v) yeast extract, 15mM maltose, 9,16 mM L-cysteine, 1,25 mM L(+)ascorbic acid, 0,77 mM KH_2_PO_4_, 3,86 mM K_2_HPO_4_, 10% (v/v) horse serum, 0,71% (v/v) iron solution [=1% (w/v) Fe(NH_4_)_2_(SO_4_) x 6H_2_O, 0,1% (w/v) 5-sulfosalicylacid]} at 37°C (Diamond, 1957).

Vaginal epithelial cells (VECs MS-74) were cultivated in 45% DMEM (Invitrogen, #31885), 45% Keratinocyte-SFM (Invitrogen, #37010022) and 10% fetal calf serum (FCS) in standard cell culture flasks (75 cm^2^) at 37°C and 5% CO_2_ in a Galaxy 48R (Eppendorf, Germany). For culture maintenance, cells were washed twice with Dulbecco’s PBS (PAA, #H15-001), digested with trypsin (Invitrogen, #25300-054) for 5 min and then inactivated with FCS. Cells were then pelletized at 755xg for 10 minutes and resuspended in fresh media and splitted 1:10 into new flasks and medium. Finally, a penicillin/streptomycin mix was added to a final concentration of 100 µg/ml to prevent bacterial contamination.

### The transcriptome of Trichomonadida

RNA-Seq reads were obtained using Illumina sequencing based on *Pentatrichomonas hominis* PhGII (NCBI, SRX2052873), *Tetratrichomonas gallinarum* M3 (NCBI, accession SRA318841), *Trichomitus batrachorum* BUB (NCBI, SRX2052874), *Trichomonas gallinae* GCB (NCBI, SRX2052872) and *Trichomonas tenax* HS-4 (NCBI, SRX2052871). RNA, which were isolated as previously described for *T. vaginalis* (Woehle et al., 2014). A quality-filtering step was applied to the reads so that the first nine nucleotide (nt) positions were rejected according to a FastQC analysis that showed low quality for the first 9 base calls. Subsequently, only reads with a minimum of 25 nt were retained. In addition, all reads containing 25% of low quality bases (25% of all bases with values ≤ Q15) identified by a self-written Perl script were rejected as well. The reads were assembled via *Trinity assembler* (vr20131110) (Grabherr et al., 2011). From all assembled contigs only the longest isoform of a candidate was selected. Open reading frames (ORFs) were identified and translated into the corresponding amino acid (aa) sequences by *getorf* from EMBOSS 6.6.0 (Rice et al., 2000) and a self-written Perl script was used to select only the longest ORF per candidate. To define an ORF, only stop codons were considered (option-find 0). Furthermore, only sequences with a minimum of 100 aa as a minimum for protein identification were used. For those sequences the best matches with *T. vaginalis* annotated genes were determined by using the BLAST program (version 2.2.28) (Altschul et al., 1997) in combination with the database TrichDB (v1.3) (Aurrecoechea et al., 2009) based on an e-value cutoff at ≤ 1e^-10^.

### Endocytic motif search

Putative Pmp and BspA protein sequences were first analyzed for the presence of a transmembrane domain (TMHMM v2.0) and only those with a minimum of one TMD were used for further examination. These sequences were then screened for the presence of endocytic motifs within the cytoplasmic tails using a custom perl script. The following search patterns were used: DxF (“D[A-Z]F”), FxDxF (“F[A-Z]D[A-Z]F”), WVxF (“WV[A-Z]F”), LLNLD (“LLNLD”), [DE]xxxL[LI] (“[DE][A-Z][A-Z][A-Z]L[LI]”), NPx[YF] (“NP[A-Z][YF]”), [FY]NPx[YF] (“[FY][A-Z]NP[A-Z][YF]”), YxxΦ (Y[A-Z][A-Z][FMVIL]"), DLYYDPM (“DLYYDPM”).

### Gene cloning and homologous expression of T. vaginalis

Candidate genes TVAG_166850, TVAG_183790, TVAG_140850, TVAG_212570 and TVAG_240680 were amplified using a proof-reading polymerase (Phusion High-Fidelity DNA Polymerase, NEB #M0530S) and cloned into our *Trichomonas* expression vectors using the SCS-Promotor (TVAG_047890) for gene expression. Thirty micrograms of plasmid DNA were used for transfection of 2.5 x 10^8^ *T. vaginalis* cells using standard electroporation (Delgadillo et al., 1997). After 4 hours of recovery, neomycine (G418) was added to a final concentration of 100 µg/ml for selection of positive transfected *T. vaginalis* cells. The correct expression of the fusion constructs was verified by specific RT PCRs (supplementary Fig. 5).

### RT PCR

For RNA isolation 50 ml of a dense grown culture was treated with Trizol™ reagent (Thermo Fisher, #15596018) according to the manufacture's guidelines. Afterwards 500 ng of RNA was applied for DNase treatment using DNase I, RNase free (Thermo Fisher, #EN0525) and cDNA was synthesized by the iScript™ Select cDNA Synthesis Kit (Biorad, #M0530S). A PCR was performed using the Phusion^®^ High-Fidelity DNA Polymerase (Biolabs, #170-8896) and the corresponding protocol. To ensure the amplification of the HA fusion constructs only, in each reaction gene specific primer were mixed with HA specific primer.

### Immunofluorescence assays

For immunofluorescent labelling 12 ml of a dense grown *T. vaginalis* culture without dead cells were centrifuged for 5 min at 900xg and RT. Supernatant was discarded and cells were gently resuspended in 500 µl fixation-buffer and incubated for 30 minutes at 37°C followed by a centrifugation at 900xg and RT for 5 minutes. Cells were gently washed in PBS and centrifuged again at same conditions. Supernatant was discarded and cells were resuspended in 100-150 µl PBS depending on size of the cell pellet. Subsequent steps were performed in a 6-Well plate (Sarstedt, #83.3925). Cell suspension was placed all-over a Poly-L-lysine (Sigma, #P4707) coated coverslide and incubated for 30 minutes. After incubation suspension was gently removed from the 6-Well slot and cells were incubated for 20 minutes in permeabilization-buffer {0,1% TritonX-100 in PBS} at RT on a 2D shaker. Alternatively, a 10 min treatment with 0.1% NP-40 was performed or 10 µg/ml digitonin was used for solubilization either for 10 or 30 min. After permeabilization cells were washed three times briefly in PBS, followed by a blocking step for 60 minutes in blocking-PBS (1% BSA, 0,25% Gelatine, 0,05% Tween20 in PBS) at RT on a 2D shaker. Blocking is followed by a brief washing step and then incubation with first antibody (Monoclonal Anti-HA, produced in mouse; Sigma, #H9658) at a concentration of 1:500 in Blocking PBS for 1 hour at RT followed by 4°C overnight. Samples were washed three times for 5, 10 and 15 minutes each before incubation with secondary antibody 1:1000 (Donkey anti-Mouse IgG, Alexa Fluor 488, Thermo Scientific, #A21202) for 2 hours at RT. After three washing steps (5, 10 and 15 min) samples were mounted with Fluorshield containing DAPI (Sigma, #F6057). Samples were stored at 4°C until imaging.

For immunofluorescence assays in the presence of vaginal epithelial cells (VECs MS-74) one day prior experiment 500 µl of VECs were given onto each slot of a 4-Well CultureSlide (BD Falcon, #354114) and incubated over night at 37°C and 5% CO_2_ in a Galaxy 48R (Eppendorf, Germany). On the next day 12 ml of a dense grown *T. vaginalis* culture were centrifuged at 900xg and RT for 5 min. The supernatant was discarded and the pellet was washed once with 500 µl PBS. After a second centrifugation step the pellet was resuspended in 100-150 µl PBS and the complete cell suspension was given onto one slot of the culture slide with preincubated VECs. After a 30 min incubation at 37°C and 5% CO_2_, fixation buffer {4% Paraformaldehyde (16%, EMS, #15710) in PBS} was added and cells were again incubated for 30 min at 37°C and 5% CO_2_. Permeabilization as well as antibody treatment was done according to the protocol above. For lysosome colocalization studies cells were first incubated with 130 nM LysoTracker™ Red DND-99 (Thermofisher, #L7528) diluted in prewarmed culture medium for 2 hours at 37°C.

### Live cell imaging

For detection of the GFP tagged fusion proteins 2 ml of a dense grown culture was centrifuged at 900 x g for 5 min and the pellet was carefully resuspended in 150 µl PBS and placed on a Poly-L-lysine (Sigma, #P4707) coated coverslide and incubated at 37°C and 5% CO_2_ in a Galaxy 48R (Eppendorf, Germany). After 1h of incubation the cells were washed once with 1 ml PBS and samples were mounted with Fluorshield containing DAPI (Sigma, #F6057) and immediately used for microscopy.

### Adhesion Assays

Adhesion assays were performed three times in triplicates for each candidate. One day prior adhesion assay 1.5×10^5^ vaginal epithelial cells (VECs MS-74) were given onto each slot of a 4-Well CultureSlide (BD Falcon, #354114) and filled up to a final volume of 1 ml with culture medium {45% DMEM (Invitrogen, #31885), 45% Keratinocyte-SFM (Invitrogen, #37010022) and 10% fetal calf serum (FCS)}. Vaginal epithelial cells were incubated over night at 37°C and 5% CO_2_ in a Galaxy 48R (Eppendorf, Germany). Preliminary to the assay *T. vaginalis* cells of a dense grown culture were incubated with 10 µM CellTracker Green CMFDA Dye (Molecular Probes, #C7025) for 30 minutes at 37°C and 5% CO_2_ in *Trichomonas* culture medium {2,22% (w/v) tryptose, 1,11% (w/v) yeast extract, 15 mM maltose, 9,16 mM L-cysteine, 1,25 mM L(+)ascorbic acid, 0,77 mM KH_2_PO_4_, 3,86 mM K_2_HPO_4_, 10% (v/v) horse serum, 0,71% (v/v) iron solution [=1% (w/v) Fe(NH_4_)_2_(SO_4_) x 6H_2_O, 0,1% (w/v) 5-sulfosalicylacid]}. Culture was spun down at 500xg for 5 minutes at RT and washed two times with *Trichomonas* culture medium. *T. vaginalis* cells were counted (TC20 Automated Cell Counter, BioRad) and for each assay infection was initiated with 5×10^4^ *T. vaginalis* cells in 500 µl *Trichomonas* culture medium and incubated at 37°C and 5°CO_2_ for 30 minutes. After incubation free swimming cells were washed off with PBS and adherent cells were fixed in PBS containing 1% Paraformaldehyde (EMS, #15710) for 30 minutes at 37°C and 5°CO_2_. Observation at microscope was performed with 10x magnification and for each assay 12 closely spaced pictures were taken. Quantitative analysis of the adherent cells was performed with ImageJ 1.48.

## Acknowledgements

We thank Peter Melzer for his help with cloning and early imaging. The funding through the DFG (CRC1208/A04) is gratefully acknowledged.

